# Photo-leucine incorporation in *Bacillus subtilis*: Identification of transporters and of a selectivity determinant in the Leu-tRNA synthetase

**DOI:** 10.1101/2025.03.07.642037

**Authors:** Robert Warneke, Christina Herzberg, Piotr Neumann, Fabian Schildhauer, Xiaoyi Xin, Ralf Ficner, Juri Rappsilber, Jörg Stülke

## Abstract

The elucidation of the intricate network of proteins is essential for our understanding of cellular function. *In vivo* crosslinking is a powerful tool for probing these interactions, with photoactivatable amino acids representing a promising next step. Here, we evaluated the application of photo-leucine in the gram-positive model organism *Bacillus subtilis*. This amino acid analog is incorporated into proteins, but high concentrations inhibit the growth of *B. subtilis*. We investigated photo-leucine homeostasis and identified the branched-chain amino acid importers BcaP and BraB as well as the bipartite exporter AzlCD as key players of its uptake and export, respectively. Additionally, we identified a previously uncharacterized exporter, AexB, a member of the "sleeping beauty" group of EamA amino acid exporters. Expression of the *aexB* gene is positively regulated by the transcription factor AerB, that is normally normally inactive. Selective pressure by photo-leucine triggered the acqusition of a point mutation in *aerB*, allowing AerB to activate *aexB* expression. Mutants highly resistant to photo-leucine carried a single amino acid exchange, Ala-494 to Thr, in the leucine-tRNA synthetase LeuS. Molecular docking analysis revealed that this mutation alters the leucine-binding pocket, in a manner that still allowed binding of leucine, but not photo-leucine. While high photo-leucine incorporation is beneficial for effective crosslinking, our results suggest that *B. subtilis* intrinsically limits its incorporation to maintain cellular tolerance. Thus, evolutionary processes provide adaptive mechanisms to adjust amino acid analogue incorporation to tolerable levels, making photo amino acids a candidate for life cell investigation of protein struture and interactions.

## INTRODUCTION

Life depends on functional interactions between biological macromolecules which must be specific with respect to the partner and with respect to time and space in the cell. Among these interactions, protein-protein interactions stand out as both the most diverse and most abundant. While the sequencing of entire genomes offers a comprehensive list of cellular components, it does not immediately unveil the specific functions of these proteins or the mechanisms by which they assemble into molecular machines and functional networks that govern cellular behavior (1, 2). The identification and structural analysis of these interactions as well as the understanding of their functions are the subject of intensive research and require a variety of methods with their strengths and weaknesses each.

Conventional methods for the identification of protein-protein interactions often rely on affnitiy-based techniques like co-immunoprecipitation, which encounter challenges when utilizing cell lysates. During the lysis procedure, false-positive interactions can occur due to the loss of spatial organization and weakly interacting proteins are often lost during the subsequent washing procedure. Alternatively, two hybrid screens are also popular. In these *in vivo* experiments, the interaction of potential partners results in a simple readout such as a growth, the induction of a reporter enzyme or fluorescence. For two-hybrid experiments, the partner proteins are typically expressed to high levels irrespective of their native timing and level of expression, and also not taking into account potential different localization. Typically, each of these approaches identifies only about one third of the actual interactions (3). To overcome all these limitations, crosslinking has emerged as a powerful method to covalently fix interactions *in vivo*, which offers a tool to detect protein-protein interactions even in challenging conditions. The classical chemical crosslinking utilizes reactive bifunctional reagents targeting free amino groups to generate covalently links proteins (4). This method has been used to study the complete *in vivo* interactome of living bacterial cells such as *Mycoplasma pneumoniae* or *Bacillus subtilis* (5, 6). It does not only provide information about interacting proteins but gives also structural information. The structural information derived from crosslinking can even be used to obtain structure models of protein complexes and the crosslinking information can be integrated into complex models predicted using artificial intelligence (6, 7, 8). This approach incorporates experimental distance restraint information, enhancing the ability of the AI algorithm to predict challenging protein structures, especially for those undergoing conformational changes or with limited homologous sequences. Despite significant success and the detection of numerous previously unknown interactions, the method has drawbacks, such as potential non-specific interactions and perturbation of the native state of proteins during the fixation process (9).

Photo-crosslinking has the potential to significantly advance *in vivo* crosslinking studies by introducing of a photo-activatable group into a protein, thus allowing the generation of highly reactive intermediates *in situ* through irradiation of an inert precursor (10, 11, 12). Due to the short lifetime of the excited intermediates, the crosslinking occurs with greater specificity, providing a major advantage. Thus, the risk of non-specific interactions is minimized, increasing the chance to detect genuine protein-protein interactions (13, 14). Beyond its precision, photo-crosslinking offers additional advantages (15). Recently, photo-crosslinking with photo-leucine was used to improve the accuracy of protein structure predictions through the introduction of AlphaLink, a modified version of the AlphaFold2 algorithm (16). Photo-crosslinking therefore presents intriguing possibilities, and its application in the well-studied model bacterium *B. subtilis* could offer an unparalleled view into the cell and its functions.

We have a long-standing interest in amino acid metabolism in *B. subtilis* (16, 17, 18, 19, 20, 21, 22). *B. subtilis* is able to import all proteinogenic as well as several non-proteinogenic amino acids; however, the responsible transporters have not yet been identified for all amino acids (23, 24). Many amino acids can inhibit the growth of bacteria as they may interfere with the synthesis of other amino acids due to structural similarities that result in mis-regulation or as a result of their rather high intrinsic reactivity that may lead to the generation of toxic metabolites (25). For *B. subtilis*, amino acid toxicity has been observed both for wild type strains and for specific mutants, such as mutants defective in the degradation of the particular amino acid or mutants lacking the essential second messenger cyclic-di-AMP. *B. subtilis* is able to cope with such toxic amino acids by either reducing their uptake or by activating their export or degradation. The known amino acid exporters in *B. subtilis* are very poorly expressed, and these exporters only become activated by the acquisition of mutations that result in their enhanced expression (22, 26, 27, 28). The recently discovered pleiotropic amino acid exporter AexA is the founding member of a group of exporters dubbed “sleeping beauty amino acid exporters”. The genes encoding these exporters belong to the most poorly expressed genes in *B. subtilis*. There is no known condition, under which the expression of these genes is induced, and mutations in the regulators are required to facilitate the expression of the exporter genes (22). In some cases, mutations affecting the target of a toxic amino acid may also help to overcome growth inhibition. This has been observed for the toxic amino acid serine, which inhibits threonine biosynthesis and thus selects for enhanced expression of threonine biosynthetic enzymes (29). Similarly, glutamate toxicity of a strain lacking cyclic di-AMP can be overcome by inactivating low affinity potassium channels since glutamate confers high affinity to those channels (20, 30).

In this study, we show that photo-leucine is incorporated into proteins of *B. subtilis* by the cognate translation machinery. This incorporation is limited to a low percentage of leucine levels and to clarify the reasons for this we embarked on a comprehensive exploration of the homeostasis of photo-leucine in *B. subtilis*. We observed that high concentrations of photo-leucine limit growth of the bacteria and made use of this growth inhibitory effect for the identification of uptake and export systems for photo-leucine. Moreover, we isolated mutants that are highly resistant to photo-leucine. Such mutants had an amino acid substitution in the leucyl-tRNA synthethase LeuS. Structure prediction and molecular docking suggested potential substrate promiscuity of LeuS, and gain in substrate selectivity in the LeuS variant. Although a high incorporation rate of photo-leucine is desired for effective crosslinking, these results suggest that the incorporation rate is naturally limited by the tolerance of *Bacillus subtilis* to this amino acid analogue. This work lays the ground for the application of photo-crosslinking in *B. subtilis* to further explore the intricate landscape of protein interactions within this model organism.

## RESULTS

### Photo-leucine is readily incorporated into proteins but limits growth of *B. subtilis* at high concentrations

Photo-leucine is a commercially available leucine analogon (see Fig. 1A), that attracted wide interest in the application of photo-crosslinking in a wide range of organisms (31, 32). To test whether *B. subtilis* is suitable for photo-crosslinking, we cultivated the wild type strain 168 in C-Glc minimal medium with or without the addition of 1 mM photo-leucine. We then lysed the cells and recovered the proteins from the lysates, allowing for the analysis of the photo-leucine incorporation rate. We observed that in approximately 3.5% of the leucine residues, the amino acid was replaced with photo-leucine. We aimed for a higher incorporation rate to increase crosslinking efficiency; however, we observed that high amounts of photo-leucine inhibit the growth of *B. subtilis*. Notably, while non-natural amino acids are often toxic for cells, photo-amino acids are non-toxic for mammalian cells and can partially replace the natural amino acids (32).

**Figure 1.**
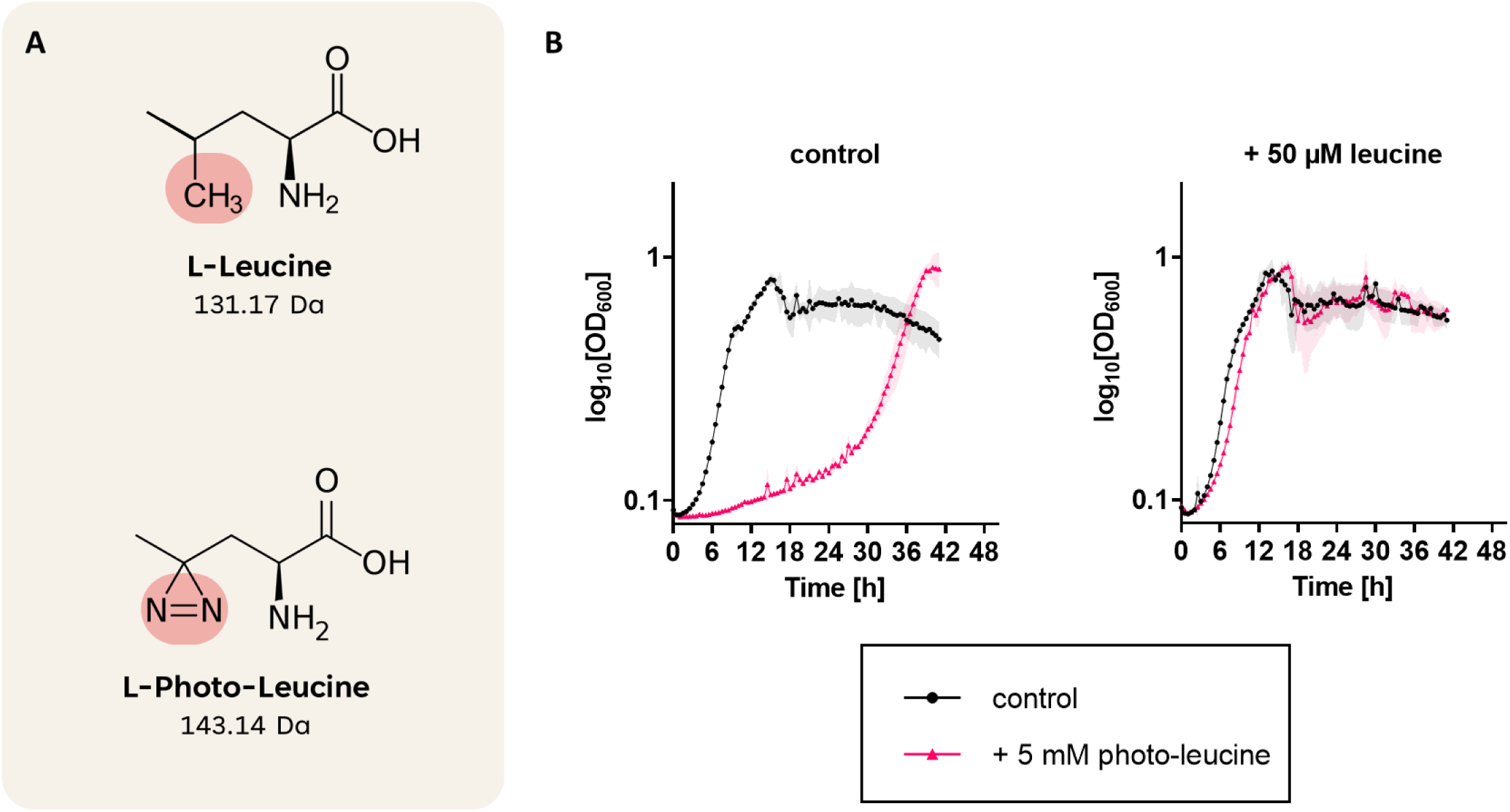
Photo-leucine inhibits growth of *B. subtilis* at high concentrations. A, The structures of L-leucine and L-photo-leucine are displayed together with their respective atomic weight. The modified side chain is highlighted. B, The wild type strain 168 was cultivated in C Glc minimal medium in presence or absence of 5 mM photo-leucine and/or 50 µM leucine.

To get a better understanding of the potential toxicity of photo-leucine, we tested the growth of the wild type strain in C Glc minimal medium with or without 5 mM photo-leucine. As shown in Fig. 1B, the strain grew in C minimal medium in the absence of photo-leucine. In contrast, the addition of photo-leucine severely inhibited growth of the strain for aproximately 24 hours before the bacteria started to grow again. It has been shown before, that in some cases toxic non-proteinogenic amino acids compete with non-toxic proteinogenic amino acids for the transport systems. The addition of the physiological amino acid can then overcome the toxicity of the toxic molecule as observed for alanine and β-alanine (22). We therefore tested growth of *B. subtilis* in the presence of both photoleucine and leucine. Indeed, the severe growth defect caused by photo-leucine was almost completelety abolished, when 50 µM L-leucine (1:100 in comparison to photo-leucine) was added to the medium (see Fig. 1B). The observation that leucine can overcome the growth inhibition caused by photo-leucine reflects the structural similarity of both molecules and suggests that they might compete with each other for the same transporter(s).

### Deletion of the L-leucine importers BcaP and BraB increases resistance to photo-leucine

Photo-leucine is an analogue of the natural amino acid L-leucine and can even be incorporated in place of it into proteins (see above, 32). In order to test whether photo-leucine may be also taken up into the cell via the same import system(s) as L-leucine, we compared the growth of the wild type strain 168 with isogenic mutant strains lacking the known leucine permeases (33) BcaP (Δ*bcaP*, BKE09460), BraB (Δ*braB*, GP4152) or both (Δ*bcaP* Δ*braB*, GP4157) in the presence of photo-leucine (see Fig. 2). In both single deletion mutants, we observed improved growth in the presence photo-leucine with the *braB* mutant performing slightly better than the *bcaP* mutant suggesting that both transporters contribute to the uptake of photo-leucine. Indeed, the *braB bcaP* double deletion mutant GP4157 showed the highest resistance against photo-leucine. These results show that photo-leucine and leucine are taken up by the same import systems, BraB and BcaP.

**Figure 2.**
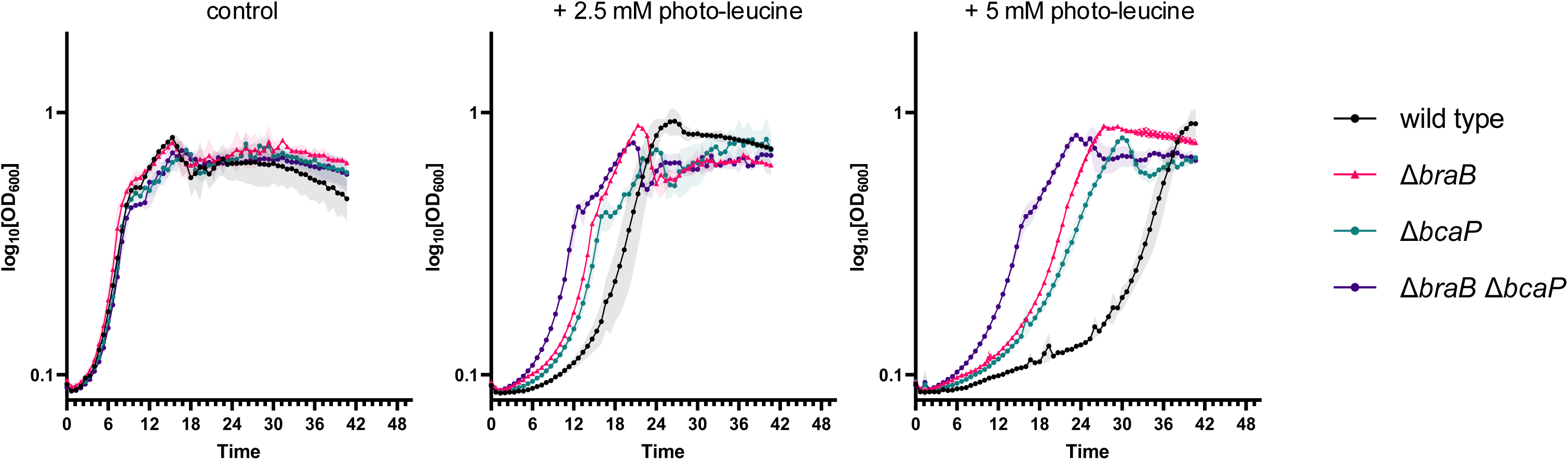
Deletion mutants lacking BcaP and BraB exhibit resistance towards photo-leucine. The growth of the wild type strain 168 (black), the Δ*braB* (GP4152) (magenta), the Δ*bcaP* (BKE09460) (turquoise), and the double mutant the Δ*braB* Δ*bcaP* (GP4157) (purple) was tested on C minimal medium in the presence of 2.5 or 5 mM photo-leucine.

### AzlCD is a major exporter for photo-leucine

To further investigate the response of *B. subtilis* to photo-leucine, we isolated suppressor mutants that are viable at otherwise growth-limiting photo-leucine concentrations. A typical feature of suppressors isolated on toxic amino acids is the inactivation of the respective importers (22, 28, 30). In order to avoid suppressors that would inactivate the known importers BcaP or BraB, we decided to work with the strain GP4157 lacking *braB* and *bcaP* for the suppressor isolation. As the strain already shows a higher resistance than the wild type (see Fig. 2), we cultivated the cells on medium containing 12 mM photo-leucine in a microplate reader. No growth was observed for several hours, but after one day growth was observed. We isolated the suppressor mutant GP4120 tolerating 12 mM of photo-leucine and subjected it to whole genome sequencing (genotypes are displayed in Fig. 3A). A mutation in the hypothetical Shine-Dalgarno sequence of the *azlB* gene encoding a transcriptional repressor of the Lrp family of the *azlB-azlC-azlD-brnQ-yrdK* operon was identified (Fig. 3B). Inactivation of AzlB leads to expression of the bipartite AzlCD complex, which exports several amino acids such as the leucine analogon 4-azaleucine, histidine, asparagine, and 2,3-diaminopropionic acid (22, 26, 27, 28). Thus, it was plausible to assume that AzlCD would also be able to export photo-leucine.

**Figure 3.**
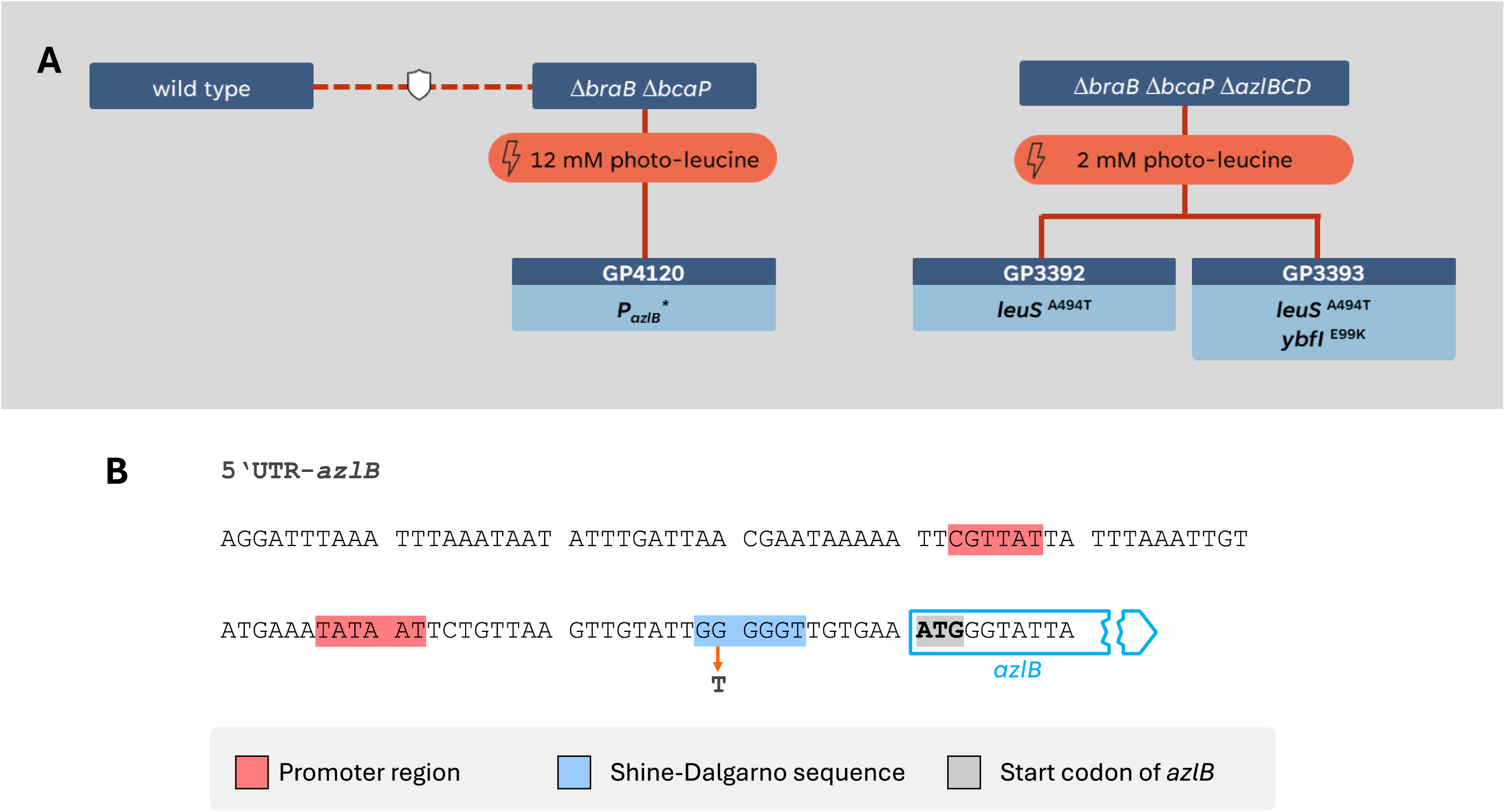
A suppressor screen in the presence of photo-leucine. A. Evolutionary trajectory of the Δ*braB* Δ*bcaP* (GP4157) Δ*braB* Δ*bcaP* Δ*azlBCD* (GP4123) strains exposed to 12 mM and 2 mM of photo-leucine, respectively. B. Location of the suppressor mutation in the Shine-Dalgarno sequence of the *azlB* gene of *B. subtilis* GP4120. The regulatory elements are labelled according to (26).

To test whether expression of AzlCD indeed conferred resistance, we tested the growth of the *azlB* (GP3600) and *azlBCD* (GP3623) deletion mutants in the presence or absence of photo-leucine (Fig. 4). Strikingly, the *azlB* mutant GP3600 which exhibits constitutive expression of the AzlCD exporter (27) showed almost no sensitivity against photo-leucine even at 5 mM photo-leucine. In contrast, the *azlBCD* mutant GP3623 was highly sensitive to photo-leucine suggesting that AzlCD is required for the resistance to photo-leucine. Moreover, this observation indicates that the very weak expression of AzlCD in the wild type strain is already sufficient to provide some basal protection of the cells against the growth-inhibiting effect of photo-leucine.

**Figure 4.**
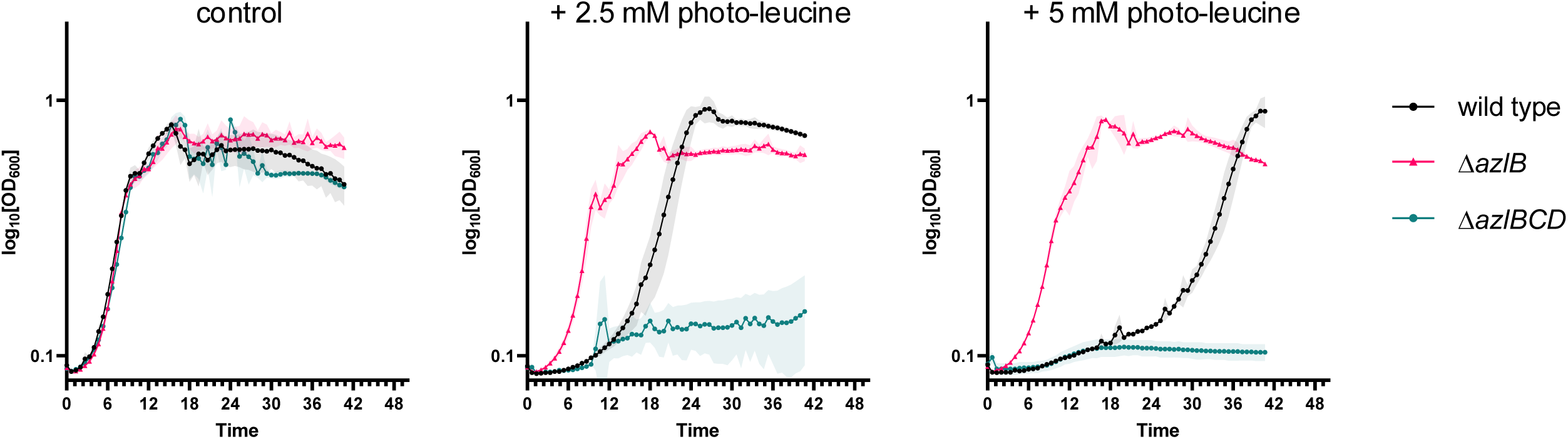
The Δ*azlB* deletion mutant exhibits strong resistance towards photo-leucine. The growth of the wild type strain 168, the Δ*azlB* (GP3600), and the Δ*azlBCD* (GP3623) was tested on C minimal medium in the presence of 2.5 or 5 mM photo-leucine.

### AerB controls the expression of the EamA family protein and photo-leucine exporter AexB

To get more insights into the growth inhibition caused by photo-leucine, we generated a *braB bcaP azlBCD* triple mutant (GP4123) lacking both the uptake and export system for photo-leucine. Probably due to the deletion of the export system AzlCD, the strain was highly succeptible to photo-leucine stress and could not grow on C Glc minimal medium supplemented with 2 mM photo-leucine (data not shown). In this strain, the most likely suppressor mutations such as the inactivation of uptake systems or activation of a major exporter are no longer possible, so we hoped that *B. subtilis* might find alternative ways to deal with the photo-leucine-induced growth inhibition. We screened for suppressors of the *braB bcaP azlBCD* triple mutant GP4123 that could tolerate 2 mM photo-leucine and isolated two independent mutants, GP3392 and GP3393. We subjected both mutants to whole genome sequencing and identified in both an identical mutation affecting the leucyl-tRNA synthetase LeuS (see below) and in one of the two suppressors (GP3393) an additional mutation in the *aerB* (previously *ybfI*) gene encoding a so far uncharacterized transcription factor (see Fig. 3).

The mutation in the *aerB* gene leads to the exchange of glutamate at position 99 to a lysine and therefore might affect the expression of another gene involved in photo-leucine resistance. AerB belongs to the AraC family of transcription factors which often control the expression of single genes or operons in their direct genomic neighborhood (34). The *aerB* gene is encoded in an operon together with the *aexB* (previously *ybfH*) gene, a member of the strongly conserved EamA family of “sleeping beauty” amino acid exporters. Recently, we characterized the founding member of this family in *B. subtilis*, AexA. The expression of the *aexA* gene is controlled by its neighbouring AraC-type transcription factor AerA (22). Members of the EamA family are typically never expressed and depend on the activation of a transcription factor by the acquisition of a mutation.

In the light of these findings, it seemed plausible that the amino acid substitution in AerB conferred DNA binding activity to the protein leading to the expression of AexB. To test these hypotheses, we fused the promoter region of the *aexB* gene to a promoterless *lacZ* reporter gene. The resulting β-galactosidase activities reflecting the promoter activity was assayed in the strains GP4436 and GP4442 that harbour the wild type *aerB* or mutant *aerB*^E99K^ alleles, respectively. The bacteria were grown in C Glc minimal medium and the β-galactosidase activities were determined (see Fig. 5A). In agreement with the global expression data (Fig. 5B; 35, 36), nearly no expression was observed in GP4442 harboring the wild type *aerB* allele. In contrast, the AerB^E99K^ protein in GP4436 allowed high promoter activity (570 units per mg of protein) as compared to the expression of genes of central carbon metabolism (37). In order to test whether the resistance against photo-leucine is caused by the activation of the *aexB* expression, we fused the *aexB* gene to a xylose inducible promotor resulting in the strain GP4424. This strain allows expression of the *aexB* gene independently from AerB. We tested the growth of GP4424 on C Glc minimal medium with photo-leucine in presence or absence of the inducer xylose (see Fig. 5C). The overexpression of AerB was both required and sufficient to confer resistance towards photo-leucine in comparison to the non-induced condition. Thus, the expression of AerB causes the AerB^E99K^-mediated resistance to photo-leucine. Based on the activity of other proteins of this family, our data suggest that AexB is a novel photo-leucine-specific amino acid exporter of the sleeping beauty family.

**Figure 5.**
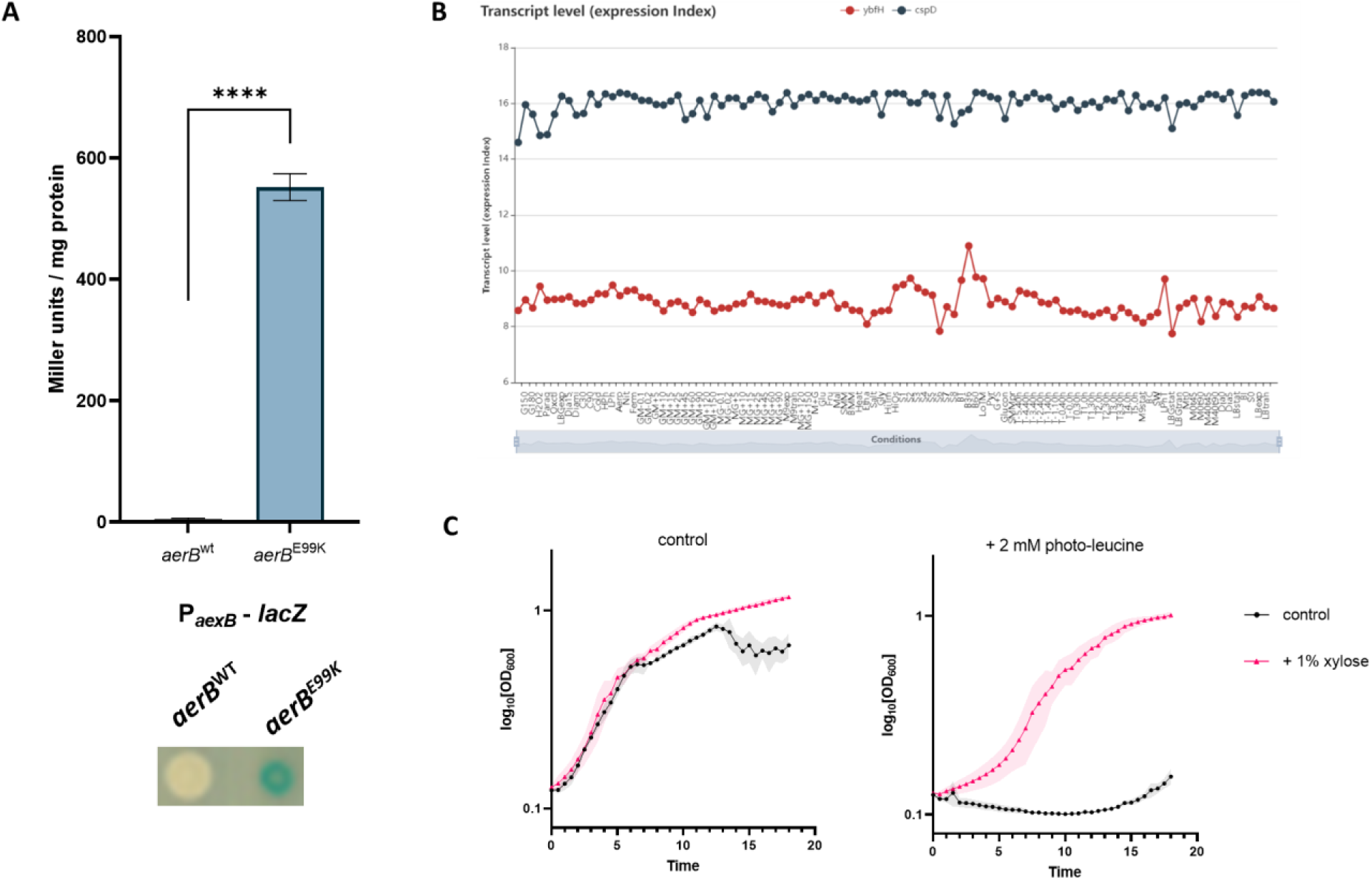
AerB controls the expression of the photo-leucine exporter AexB. A, The expression of the *aexB* promotor was monitored in isogenic wild type and the *aerB*^E99K^ strains that harbor the *aexB-lacZ* reporter gene fusion. The values for the β-galactosidase activity represent three independently grown cultures, and for each sample, enzyme activity was determined twice. B, The Expression Browser of *Subti*Wiki (36) allows a direct comparison of the expression levels of two or more genes (here *cspD* and *aexB*). The expression level of *cspD* serves as an example of a gene with high constitutive expression, while the *aexB* gene is virtually never expressed. C, The strain GP4424 that harbors a xylose-inducible *aexB* gene was cultivated in C minimal medium with 2 mM photo-leucine with or without 1 % xylose.

### An amino acid exchange in leucyl-tRNA synthetase confers high resistance against photo-leucine

As mentioned above, both suppressor mutants isolated from the *braB bcaP azlBCD* triple mutant GP4123 had an identical mutation in the *leuS* gene, which resulted in a A494T substitution in the leucyl-tRNA synthetase LeuS (see Fig. 3). According to InterPro (38), the amino acids 406-604 of LeuS form the IPR002300 amino acyl-tRNA synthetase IA domain which catalyzes the attachment of an amino acid to its cognate transfer RNA molecule in a highly specific two-step reaction. The structure of LeuS in complex with substrate analogs has been determined for several bacteria (see Fig. 6). While the neighbouring residues Gln-492 and Trp-493 are part of an alpha helix that constitutes the leucine binding pocket, the Ala-494 is facing to the backside of the this alpha helix and is thus not directly part of the substrate binding site. Since the *leuS* gene is essential (39), the options for mutations that retain the function of the protein are probably very limited. It is therefore not surprising that we identified the same mutation in LeuS in both independent suppressors. We compared the growth of the *braB bcaP azlBCD* mutant with the isogenic *leuS*^A494T^ in C Glc minimal medium supplemented with 5 mM photo-leucine. Strikingly, the mutant carying the *leuS* variant showed strong resistance against photo-leucine (see Fig. 7).

**Figure 6.**
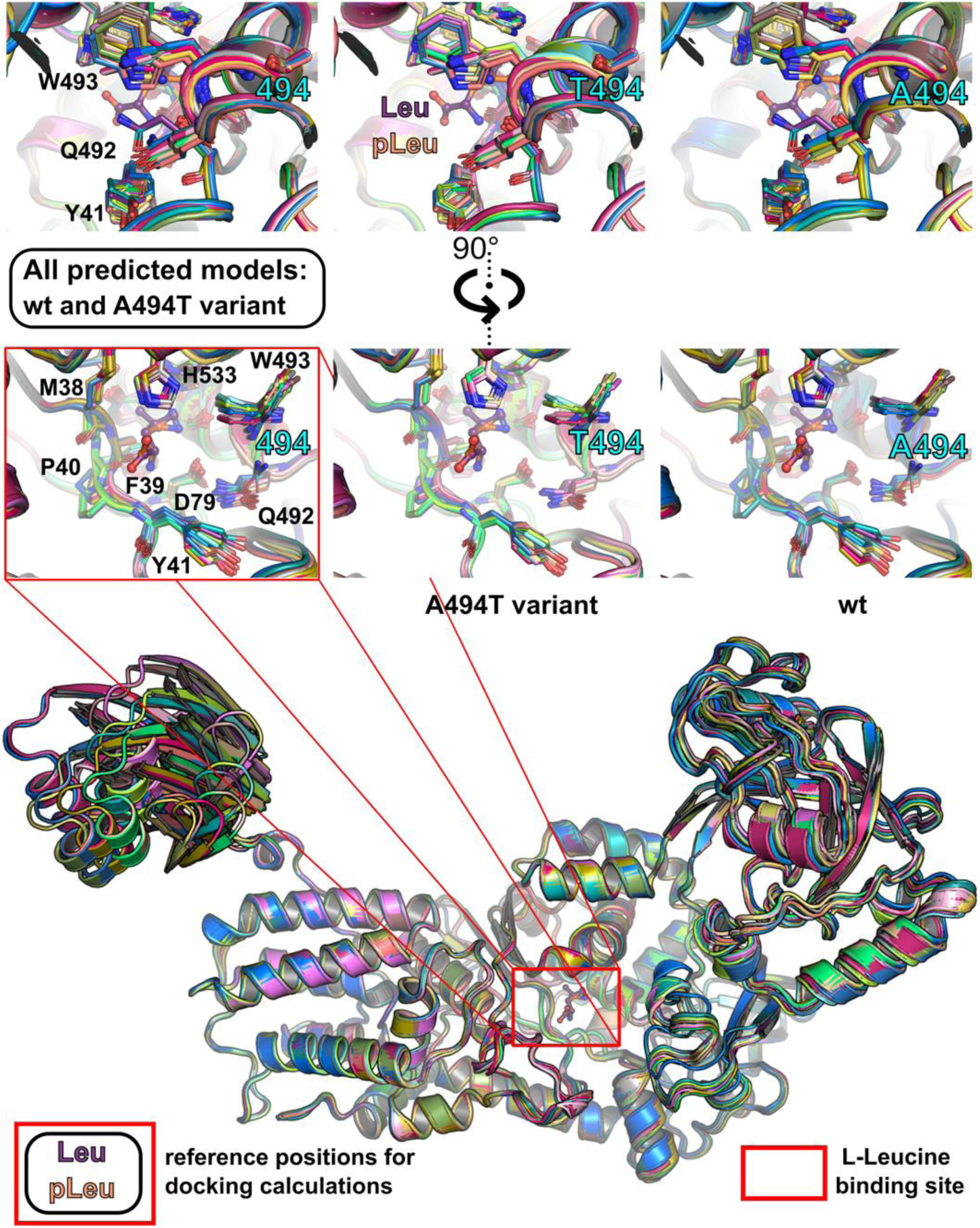
Superposition of three crystal structures (2V0C, 7NU0, 5AH5), shown as cartoon, structurally related to *Bs*LeuS. The crystallographically determined ligand molecules are shown as sticks. The reference positions of the leucine and photo-leucine molecules used for the docking experiments are shown as balls and sticks. pLeu, photo-leucine.

**Figure 7.**
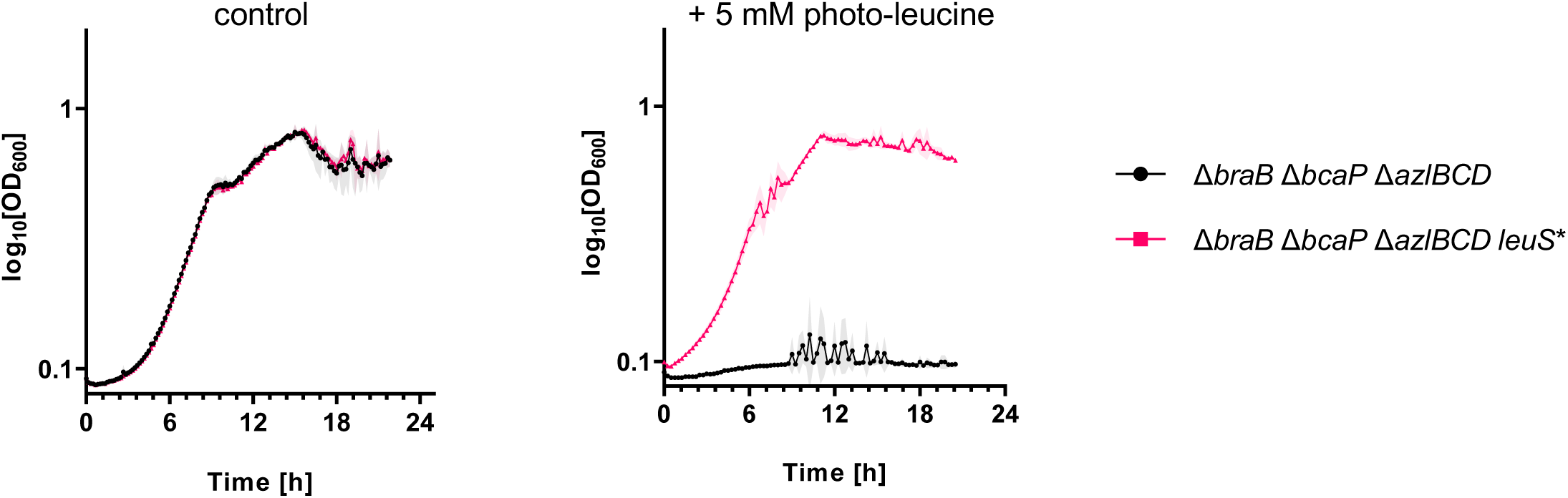
LeuS directly influences the resistance against photo-leucine. A, The growth of the Δ*braB* Δ*bcaP* Δ*azlBCD* (GP4123, black) and the isogenic strain encoding the LeuS variant (GP3392, magenta) was tested on C minimal medium in the presence of 5 mM photo-leucine.

### Molecular docking of leucine and photo-leucine to the wild type and A494T LeuS

To investigate how the A494T mutation in leucyl-tRNA synthetase (LeuS) may alter substrate specificity, we performed molecular docking using ColabFold-predicted structures and GNINA (40, 41). The structural comparison revealed that the mutation caused subtle positional shifts in residues P491-G495, leading to a reorientation of the bulky Trp-493 and Gln-492 side chains. As a consequence, these structural alterations lead to a change in the volume of the L-leucine binding pocket. Therefore, they could be responsible for the physiologically observed discrimination against photo-leucine observed in the A494T variant, which, in contrast to the wild type LeuS, does not accept L-photo-leucine as a substrate.

Docking simulations using the Vinardo scoring function (42) confirmed that wild-type LeuS could accommodate both leucine and photo-leucine, whereas the A494T variant did not allow binding of photo-leucine due to steric hindrance and altered side-chain interactions. Based on the atomic structure prediction, incorporation of two additional atoms near the Cβ atom of alanine 494 (A494T mutation) leads to structural alterations that result in a slight repositioning of residues 491-495 and a significant change in the W493 side-chain conformation (see Fig. 8). It is tempting to speculate, that the resulting repositioning of the bulky tryptophan side chain may be directly responsible for the gain in substrate specificity of the LeuS A494T variant. Thus, this mutation leads to a significant modulation of the narrow hydrophobic pocket suited for binding of a leucine side chain (see Fig. 8). The pocket becomes wider and more polar. The mutation thus increased selectivity for leucine, preventing misincorporation of the analog while maintaining the essential enzymatic function. These findings highlight how minimal structural modifications can fine-tune substrate discrimination, offering insights into the design of more selective tRNA synthetases for amino acid analog applications.

**Figure 8.**
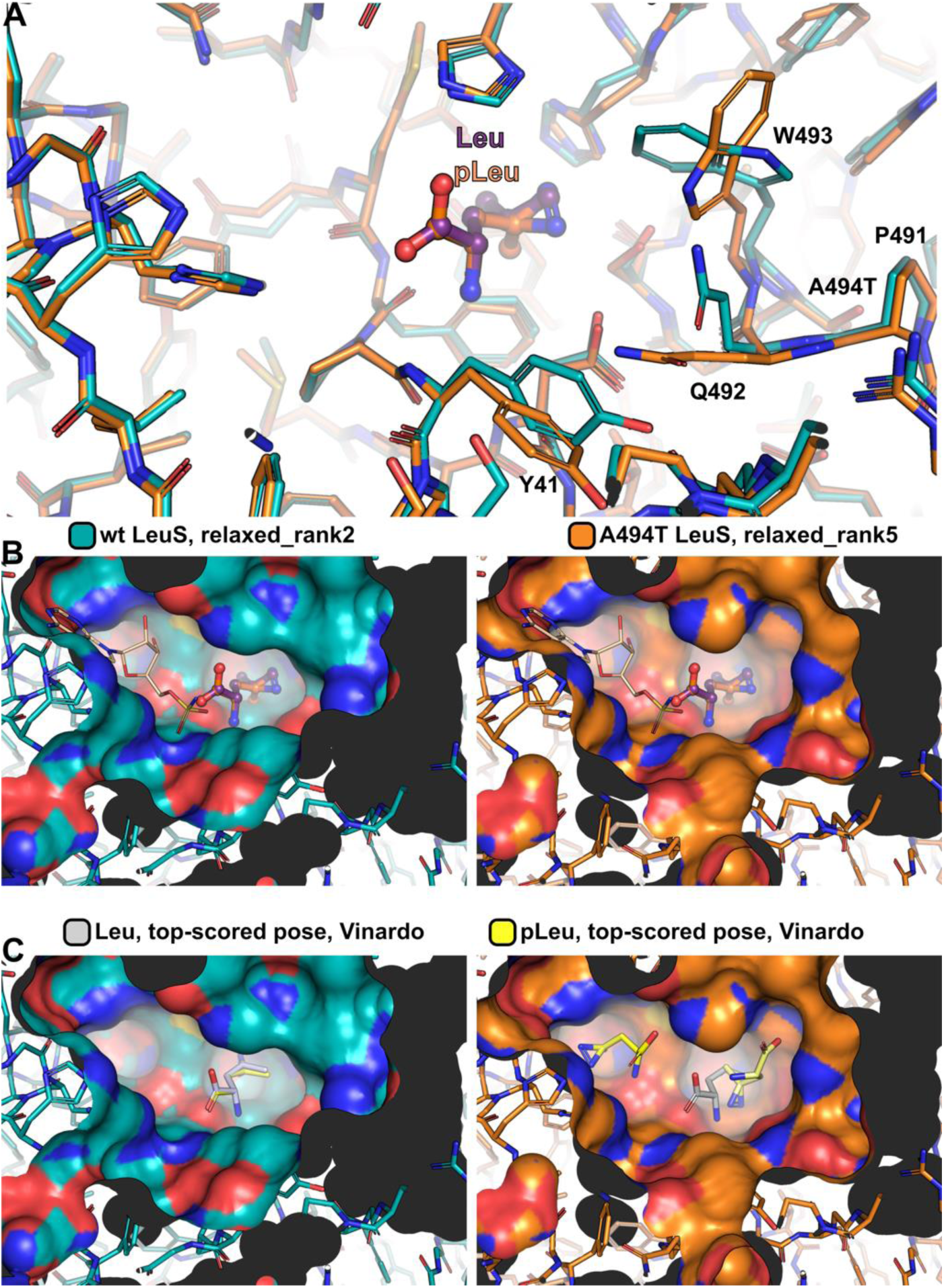
Structural explanation of the gain in substrate specificity. A, AlphaFold predicted atomic models showing the structural differences caused by the mutation from alanine to threonine at position 494 (A494T). The reference positions of the Leu and photo-leucine ligands are shown as balls and sticks. B, Surface representation (coloured by atom) of the two atomic models shown in A. The adenosine binding pocket is marked by the sulfamoyl analogue of leucyl adenylate (wheat sticks) derived from the 2V0C structure. The orientation corresponds to that of A. C, Docking results of Leu and photo-leucine ligands to selected wt and A494T models (Supplemental Table S1). The orientation corresponds to that of A.

## DISCUSSION

In this study, we investigated the incorporation of and the cellular response to the photo-amino acid photo-leucine in *B. subtilis* (see Fig. 9 for an overview). This amino acid analog is thought not to interfere with the physiology of human cells (32). In contrast, our results demonstrate that photo-leucine can be readily incorporated into proteins; however, it limits growth of *B. subtilis* at high concentrations, leading to growth inhibition. The incorporation photo-leucine into proteins is the prerequisite for its application in crosslinking experiments to identify protein-protein interactions. This incorporation requires addition of photo-leucine to the leucine-specific tRNA molecules by the leucyl-tRNA synthetase, and subsequently its addition to nascent peptide chains in the ribosome.

**Figure 9.**
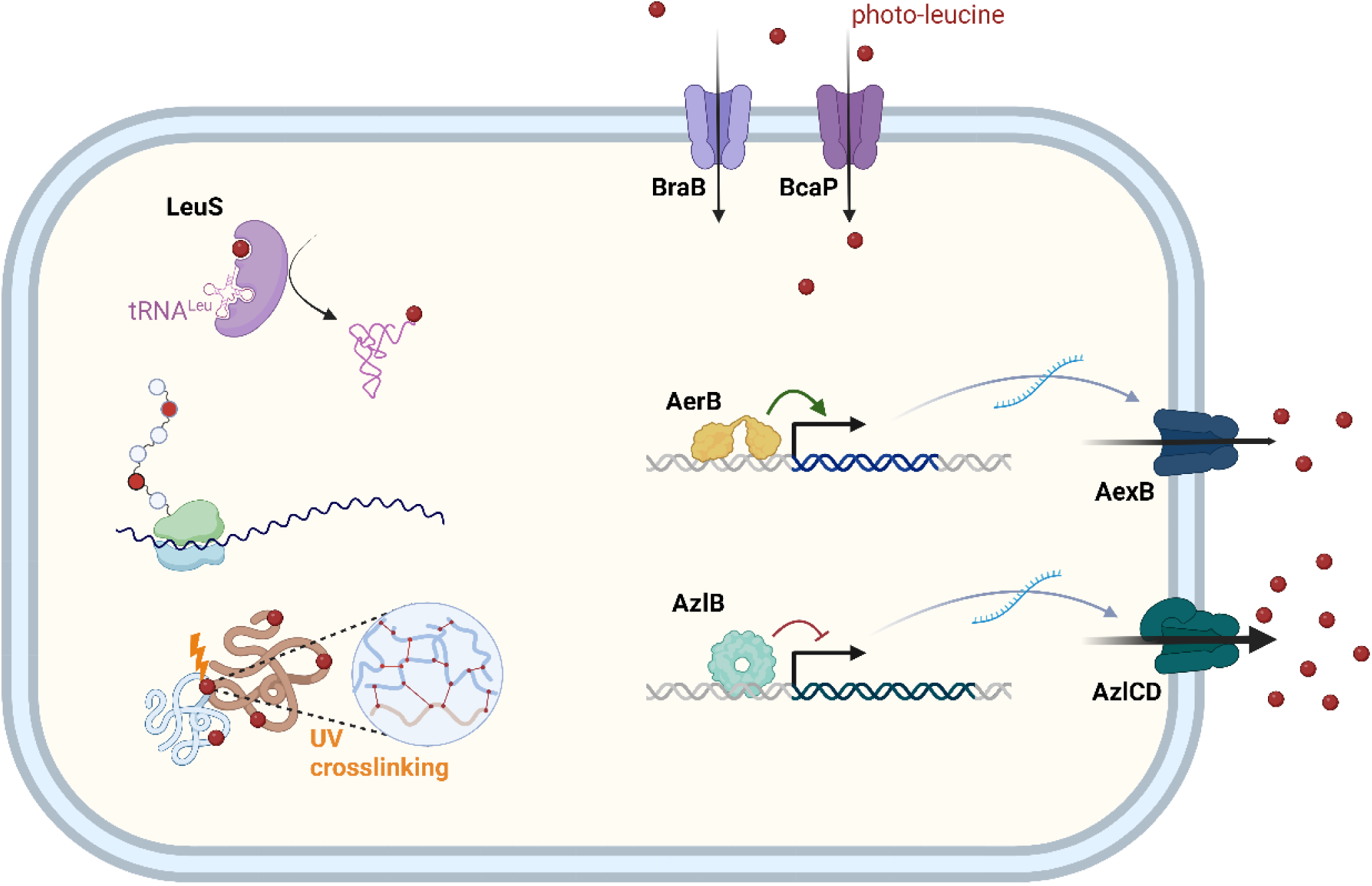
Schematic representation of photo-leucine homeostasis in *B. subtilis*.

Through a series of experiments, we identified import and export systems involved in the uptake and efflux of photo-leucine, shedding light on the mechanisms underlying its homeostasis. Furthermore, our investigation into the molecular mechanism of photo-leucine incorporation revealed insights into the substrate specificity of leucyl-tRNA synthetase LeuS, a key enzyme involved in protein synthesis. Through molecular docking simulations and structural analysis, we elucidated how a specific amino acid substitution in LeuS confers altered substrate specificity, leading to resistance against photo-leucine.

Photo-leucine is a molecule that is not occuring in nature suggesting that the leucyl-tRNA synthetase LeuS does not need to discriminate between leucine and photo-leucine. This results in the misincorporation of photo-leucine into proteins at places where normally leucine is incorporated. Such incorporation limits bacterial growth, possibly because of stable aggregates form due to basal activation by light in the laboratory. Alternatively, the protein molecules carrying photo-leucine might misfold and thus lack the required activity. The high selectivity of the A494T variant of the enzyme prevents the misincorporation of photo-leucine. Of course, this also interferes then with the possibility to get intended protein crosslinks in interaction studies.

Amino acid uptake in *B. subtilis* typically involves multiple transporters with differing affinities and selectivities. This is also the case for the non-natural amino acid photo-leucine. Two permeases, BraB and BcaP, contribute to the uptake of this artificial amino acid. BraB seems to be specific for branched-chain amino acids, including isoleucine, valine, and photo-leucine (33, this work), whereas BcaP is a highly promiscous amino acid transporter, as it contributes to the uptake of amino acids as diverse as isoleucine, valine, threonine, serine, asparagine, diaminopropionic acid, and photo-leucine (22, 28, 30, 33, this work). Based on the growth of the *bcaP* and *braB* mutants, BraB seems to play a major role in photo-leucine transport.

As observed for other amino acids or amino acid analogs, the efficient export of the amino acids plays a major role in the adaptation to these growth-inhibiting molecules. As amino acids have to be taken up or to be synthesized by the cell, a lot of energy is invested for their acquisition. This may explain why the genes encoding amino acid exporters are often only very poorly expressed in *B. subtilis*. This is the case for the well-characterized exporters AzlCD and AexA (26, 27), but also for the photo-leucine exporter AexB discovered in this work. Their corresponding genes belong to the most poorly expressed genes in *B. subtilis* (35, 36). The bipartite exporter AzlCD seems to be rather unspecific since its activation can confer resistance to a wide range of amino acids, including histidine, asparagine and diaminopropionic acid as well as to the leucine analogs 4-azaleucine and photo-leucine (22, 26, 27, 28). In this respect, the amino acid exporter AzlCD resembles the amino acid importers which also often transport multiple substrates. In addition to AzlCD, we also identified AexB, a second transporter which can export photo-leucine as soon as its expression is switched on. As for the *azlCD* and *aexA* genes, there is no known condition under which the *aexB* gene is naturally expressed.

This study also identified a transcription factor, AerB, that controls the expression of the photo-leucine exporter AexB, highlighting the intricate regulatory networks governing amino acid homeostasis in *B. subtilis*. AerB is an inactive transcription factor that can only be activated by the acquisition of particular point mutations that then allow *aexB* expression. AexB is therefore the second characterized member of the group of Sleeping Beauty amino acid exporters. The activation of AerB by point mutations is reminiscent of other transcription factors that can be activated or become independent from cofactors such as the *E. coli* cAMP receptor protein Crp, the *Listeria monocytogenes* virulence transcription factor PrfA, and the *B. subtilis* cryptic activator AerA (22, 43, 44, 45).

To use photo-leucine and potentially other photo-amino acids for *in vivo* protein crosslinking studies, there seems to be a trade-off between an intended high incorporation rate and the biological tolerance to the amino acid analog. The identification of a variant of the leucyl-tRNA synthetase with increased selectivity suggests a way to construct enzymes that provide an optimal balance between the two parameters. LeuS variants carrying mutations close to the amino acid binding site that are sufficiently selective for leucine to allow growth but still give rise to tRNA molecules charged with photo-leucine to faciilitate crosslinking of interacting proteins would allow the application of photo-leucine for *in vivo* crosslinking in *B. subtilis*. Such leucyl-tRNA synthetases could not only be used in *B. subtilis*, but in a wide variety of organisms. Moreover, such a strategy might also work for the use of other photo-amino acids.

## MATERIALS AND METHODS

### Strains, media and growth conditions

*E. coli* DH5α (46) was used for cloning. All *B. subtilis* strains used in this study are derivatives of the laboratory strain 168. They are listed in Table 1. *B. subtilis* was grown in Luria-Bertani (LB) or in sporulation (SP) medium (46, 47). Growth experiments were performed with C Glc minimal medium that contains glucose and ammonium as the sources of carbon and nitrogen, respectively (48). The media were supplemented with ampicillin (100 µg/ml), kanamycin (10 µg/ml), chloramphenicol (5 µg/ml), spectinomycin (150 µg/ml), or erythromycin and lincomycin (2 and 25 µg/ml, respectively) if required.

**Table 1.** Bacterial strains used in this study.

|  | Genotype | Reference |
| --- | --- | --- |
| 168 | *trpC2* | Lab collection |
| BKE09460 | *trpC2 ΔbcaP::ermC* | 39 |
| GP3392 | *trpC2 ΔbcaP::ermC ΔbraB::aphA3 ΔazlBCD::spc leuS<sup>A494T</sup>* | Suppressor of GP4123 on C<br>Glc + 2 mM photo-leucine |
| GP3393 | *trpC2 ΔbcaP::ermC ΔbraB::aphA3 ΔazlBCD::spc leuS<sup>A494T</sup> aerB<sup>E99K</sup>* | Suppressor of GP4123 on C<br>Glc + 2 mM photo-leucine |
| GP3600 | *trpC2 ΔazlB::spc* | 27 |
| GP3623 | *trpC2 ΔazlBCD::spc* | 27 |
| GP4152 | *trpC2 ΔbraB::aphA3* | This work |
| GP4157 | *trpC2 ΔbcaP::ermC ΔbraB::aphA3* | BKE09460 → GP4152 |
| GP4120 | *trpC2 ΔbcaP::ermC ΔbraB::aphA3 5'UTR<sup>azlB</sup><sub>(GTTGTATTGG→GTTGTATTGT)</sub>* | Suppressor of GP4157 on C<br>Glc + 12 mM photo-leucine |
| GP4123 | *trpC2 ΔbcaP::ermC ΔbraB::aphA3 ΔazlBCD::spc* | GP3623 → GP4157 |
| GP4424 | *trpC xkdE::P<sub>xyI</sub>-aexB-ermC* | pGP4003 → 168 |
| GP4436 | *trpC2 ΔbraB::aphA3 ΔbcaP::ermC ΔazlBCD::spc leuS<sup>A494T</sup> aerB<sup>E99K</sup> amyE::P<sub>aerB</sub>-lacZ-cat* | pGP4408 → GP3393 |
| GP4442 | *trpC2 ΔbraB::aphA3 ΔbcaP::ermC ΔazlBCD::spc leuS<sup>A494T</sup> amyE::P<sub>aerB</sub>-lacZ-cat* | pGP4408 → GP3392 |

### DNA manipulation and transformation

All commercially available restriction enzymes, T4 DNA ligase and DNA polymerases were used as recommended by the manufacturers. DNA fragments were purified using the QIAquick PCR Purification Kit (Qiagen, Hilden, Germany). DNA sequences were determined by the dideoxy chain termination method (46). Standard procedures were used to transform *E. coli* (46), and transformants were selected on LB plates containing ampicillin (100 µg/ml). *B. subtilis* was transformed with plasmid or chromosomal DNA according to the two-step protocol described previously (47). Transformants were selected on SP plates containing chloramphenicol (Cm 5 µg/ml), kanamycin (Km 10 µg/ml), spectinomycin (Spc 150 µg/ml), or erythromycin plus lincomycin (Em 2 µg/ml and Lin 25 µg/ml).

### Construction of mutant strains by allelic replacement

Deletion of genes was achieved by transformation of *B. subtilis* 168 with a PCR product constructed using oligonucleotides to amplify DNA fragments flanking the gene of interest and intervening resistance cassette specifying antibiotic resistance as described previously (49). The integrity of the regions flanking the integrated resistance cassette was verified by sequencing PCR products of about 1,100 bp amplified from chromosomal DNA of the resulting mutant strains.

### Phenotypic analysis

In *B. subtilis*, amylase activity was detected after growth on plates containing nutrient broth (7.5 g/l), 17 g Bacto agar/l (Difco) and 5 g hydrolyzed starch/l (Connaught). Starch degradation was detected by sublimating iodine onto the plates. Quantitative studies of *lacZ* expression in *B. subtilis* were performed as follows: cells were grown in C-Glc minimal medium. Cells were harvested at an OD_600_ of 0.5 to 0.8. β-Galactosidase specific activities were determined with cell extracts obtained by lysozyme treatment as described previously (47). One unit of β-galactosidase is defined as the amount of enzyme which produces 1 nmol of o-nitrophenol per min at 28°C.

### Genome sequencing

Chromosomal DNA from *B. subtilis* was isolated using the peqGOLD Bacterial DNA Kit (Peqlab, Erlangen, Germany). To identify the mutations in the suppressor mutants, their genomic DNA was subjected to whole-genome sequencing. Briefly, the reads were mapped on the reference genome of *B. subtilis* 168 (GenBank accession number: NC_000964) (50). Mapping of the reads was performed using the Geneious software package (Biomatters Ltd., New Zealand) (51). Frequently occurring hitchhiker mutations (52) and silent mutations were omitted from the screen. The resulting genome sequences were compared to that of our in-house wild type strain. Single nucleotide polymorphisms were considered as significant when the total coverage depth exceeded 25 reads with a variant frequency of ≥90%. All identified mutations were verified by PCR amplification and Sanger sequencing.

### Plasmid constructions

The plasmid pAC7 (53) was used to construct a translational fusion of the potential *aerB-aexB* operon promoter region to the promoterless *lacZ* gene. For this purpose, the promoter region was amplified using oligonucleotides that attached *EcoR*I and *BamH*I restriction to the ends of the product. The fragment was cloned between the *EcoR*I and *BamH*I sites of the plasmid. The resulting plasmid was pGP4408. To allow for the ectopic expression of the *aexB* gene, we constructed the plasmid pGP4003. The *aexB* gene was amplified using oligonucleotides that added *BamH*I and *Xba*I sites to the ends of the fragment and cloned into the integrative expression vector pGP886 (54) that was linearized using the same restriction enzymes.

### Analysis of photo-leucine incorporation

An equal amount of proteins isolated from *B. subtilis* 168 cultures grown in C-Glc minimal medium with or without the addition of 1 mM photo-leucine were loaded onto NuPAGE™ 4–12% Bis-Tris protein gels (Invitrogen™) and run for 10 min at 200 V. The gel was stained with Coomassie Blue to visualize the area containing proteins, the stained gel part was cut and destined in water. The proteins were in gel digested with trypsin (55). After peptide extraction from the gel, the peptides were desalted using C18-StageTips (56). Sample preparation and subsequent analyses were done in triplicate.

Peptides from in-gel digestion were acquired in a Q Exactive HF mass spectrometer (Thermo Fisher Scientific, Bremen, Germany) coupled to an Ultimate 3000 RSLC nano system (Dionex, Thermo Fisher Scientific, Sunnyvale, USA), operated under Tune 2.9, SII for Xcalibur 1.4 and Xcalibur 4.1. 0.1% (v/v) formic acid and 80% (v/v) ACN, 0.1% (v/v) formic acid served as mobile phases A and B, respectively. Samples were loaded in 1.6% acetonitrile, 0.1% formic acid on an Easy-Spray column (C18, 50 cm, 75 μm ID, 2 μm particle size, 100 Å pore size) operated at 45°C and running with 300 nl/min flow. Peptides were eluted with the following gradient: 2 to 4% buffer B in 1 min, 4 to 6% B in 2 min, 6 to 37.5% B in 72 min, 37.5 to 42.5% in 5 min, 42.5% to 50% in 6 min, 50% to 90% in 3 min and hold at 90% for 7.5 min followed by 90 to 2%B in 23.5 min. For the mass spectrometer, the following settings were used: MS1 scans resolution 120,000, AGC target 3 × 106, maximum injection time 50 ms, scan range from 350 to 1600 m/z. The ten most abundant precursor ions with z = 2–6, passing the peptide match filter (“preferred”) were selected for HCD (higher-energy collisional dissociation) fragmentation employing stepped normalized collision energies (29 ± 2). The quadrupole isolation window was set to 1.6 m/z. The minimum AGC target was 2.50 × 104, and the maximum injection time was 50 ms. Fragment ion scans were recorded with a resolution of 15,000, AGC target set to 1 × 105, scanning with a fixed first mass of 100 m/z. Dynamic exclusion was enabled for 30 s after a single count and included isotopes. Each LC-MS acquisition took 120 min. Raw data from LC-MS runs were searched using MaxQuant 1.6.17..

Files from raw data obtained from linear identification of proteins were pre-processed using MaxQuant 1.6.17.. Default settings with minor changes were used: three allowed missed cleavages; up to four variable modifications per peptide including oxidation on Met, acetylation on protein N-terminal as well as the modification of L-leucine to L-photo-leucine. Carbamidomethylation on Cys was set as fixed. The search ‘matching between runs’ feature was enabled with default settings. The full *B. subtilis* strain 168 proteome with 4,191 (Swiss-Prot) proteins was used. For protein quantification, two or more peptides using the iBAQ approach were applied. For an open modification search MSFragger (version 4.0) with full *B. subtilis* strain 168 proteome with 4,191 (Swiss-Prot) proteins was used. A mass shift at leucine sites of 11.9 Da was matched to a substitution of L-leucine to L-photo-leucine.

### Atomic model generation with ColabFold

The local installation of ColabFold (40), which is compatible with AlphaFold 2.3.2, was used to predict atomic models of the LeuS wild type (wt) and its A494T variant (A494T). For both modeling jobs, the default options were kept except for the following two adjustments. The first was the number of “prediction recycles", which was set to 10 (default 0) and the second was the activation of OpenMM/Amber for structure relaxation (default False). The first setting can improve the quality of prediction, while the second can improve the quality of side chains, both at the cost of a longer runtime. A total of 5 unrelaxed and 5 relaxed atomic models of wt and A494T *Bs* LeuS were calculated and subsequently analyzed. The models were scored based on the predicted local distance difference test (pLDDT) and predicted template modeling score (pTM) scores. The pLDDT is a measure of local accuracy and corresponds to the model’s predicted per residue scores on the local distance difference test (lDDT-Cα). The pTM measures the structural congruence between two folded protein structures. The reported values of the pLDDT and pTM scores were between 91.4/0.831 and 92.2/0.849 for the wild type protein and 91.5/0.828 and 92.4/0.849 for the A494T variant models. The reported scores imply both a very high accuracy (pLDDT > 90) and confidence (pTM > 0.8) of the predicted models. These wild type and mutant models were overlaid with the model revealing the highest pLDDT/pTM scores (A494T rank 1) and showed a very high structural similarity (see Fig. 6), as judged by the calculated RMSDs between the common Cα positions. The calculated RMSDs between the ten A494T models ranged from 0.079 Å (for 778 positions) to 0.333 Å (for 644 positions) and were slightly lower than those between the wild type models and the A494T rank 1 reference model (from 0.312 Å for 628 positions to 0.600 Å for 617 positions).

### Molecular docking with GNINA, assessment of pose and scoring (ranking) accuracy

Docking of L-leucine and L-photo-leucine molecules was performed with the molecular docking program GNINA (41). The leucine binding site was deduced based on the overlay with the structurally related crystal structures of the *Neisseria gonorrhoeae* Leucine-tRNA ligase (PDB id: 7NU0, identity 46%, similarity 81.5%, complex with L-leucinol), *Agrobacterium radiobacter* K84 Leucine-tRNA ligase (PDB id: 5AH5, identity 40%, similarity 74.8%, complex with 5’-O-(L-leucylsulfamoyl)adenosine), and *Thermus thermophilus* Leucyl-tRNA synthase (PDB id: 2V0C, identity 50%, similarity 90.2%, complex with sulphamoyl analogue of leucyl-adenylate). The superposition of these three crystal structures, performed using the active site regions, revealed that the leucine moieties of the bound ligands are almost identically positioned. Therefore, we used them to determine the reference positions of the leucine and photo-leucine molecules that we intended to use for the docking experiments. The reason for this step was to simplify the analysis of the docking results, which we wanted to perform by calculating the root mean square deviation (RMSD) between the docked poses and the reference position. The calculation of the RMSD should allow a fast and accurate quantification of the docking experiments, *e.g.*: a small RMSD indicates that the docked pose is positioned similarly to the reference molecule and *vice versa*. In addition, these three crystal structures were also used to evaluate the predictive power of the *in silico* docking approach by redocking experiments of the leucine molecule (real substrate) to the known binding pocket of each of these structures. Here, low (> 1.0 Å) RMSDs would indicate that GNINA was able to properly predict the position of leucine within the binding site. This evaluation of pose accuracy was combined with the evaluation of scoring accuracy by testing two commonly used scoring functions: Vinardo and Vina. These two scoring functions could provide different results and one of them could be more suited for the current docking experiments than the other. To this end, two sets of docking calculations were performed, using the default settings in each case, except for exhaustiveness, which was changed from its default value of 8 to 16, 32 and 64. Exhaustiveness changes the amount of computational effort during a docking experiment and a higher value (> 8) could lead to a more consistent docking result. In summary, GNINA pose accuracy and scoring (ranking) evaluations were based on comparing the top-scoring docked leucine position with the leucine reference position (crystallographic pose) by calculating the RMSD using the obrms program from the OpenBabel distribution (version 3.1.0). These docking experiments revealed that GNINA in combination with the Vinardo scoring function can properly predict the position of leucine in the known active site. Consequently, the same docking and scoring protocol was used to model the binding modes of leucine and photo-leucine in the ColabFold/AlphaFold predicted models of wild type and A494T LeuS. The reference position of photo-leucine was modeled based on the overlay with the crystallographic position of leucine (PDB id: 2V0C), using the pair fitting wizard in PyMol (Open-Source build, version 2.6.0, https://pymol.org). The orientation of the 3*H*-diazirine (>CN2) ring was arbitrarily placed, thus slightly larger RMSD values (∼ 1.5 Å) could still indicate an almost identical position when comparing the docked pose with the reference molecule. Based on visual inspection, the following RMSD thresholds were applied to distinguish whether the predicted ligand position is similar to the crystallographic pose or not (leucine – 1.5 Å, photo-leucine 2.0 Å). An RMSD greater than 5.0 Å was interpreted as no binding event.

## Acknowledgements

This work was supported by the Deutsche Forschungsgemeinschaft via the Priority Program SPP1879 (project Stu 214/16-2) (to J.S.) and Germanýs Excellence Strategy – EXC 2008 – 390540038 – UniSysCat (to J.R.).

## Author contributions

RW, JR, RF, and JS conceived and designed the experiments. RW, CH, FS, and XX performed the experiments. PN computed the structures. RW, CH, PN, FS, RF, JR and JS analyzed and discussed the data; RW, FS and JS wrote the manuscript. All authors edited and approved the manuscript.

