## Supplementary material for "Photo-leucine incorporation in *Bacillus subtilis*: Identification of transporters and of a selectivity determinant in the Leu-tRNA synthetase": Table S1

**Table S1. Molecular docking results of leucine and photo-leucine ligands docked to wt and A494T variant of *Bs*LeuS. Models marked with # are depicted in Figure 8. Ex, exhaustiveness.**

| **Ex.** | **WT ColabFold model** | **leucine RMSD [Å]** | **photo-leucine RMSD [Å]** | **Ex.** | **A494T ColabFold model** | **leucine RMSD [Å]** | **photo-leucine RMSD [Å]** |
|---|---|---|---|---|---|---|---|
| **8** | relaxed_rank1 | 1.442 | 3.538 | 8 | relaxed_rank1 | 3.247 | 2.061 |
| **8** | relaxed_rank2 | 0.743 | 7.918 | 8 | relaxed_rank2 | 0.717 | 1.773 |
| **8** | relaxed_rank3 | 7.622 | 2.986 | 8 | relaxed_rank3 | 2.285 | 6.598 |
| **8** | relaxed_rank4 | 3.839 | 3.522 | 8 | relaxed_rank4 | 5.978 | 3.587 |
| **8** | relaxed_rank5 | 2.687 | 1.635 | 8 | not-relaxed_rank1 | 0.978 | 7.593 |
| **8** | not-relaxed_rank2 | 0.7 | 7.633 | 8 | not-relaxed_rank2 | 0.72 | 1.789 |
| **8** | not-relaxed_rank3 | 1.22 | 7.763 | 8 | not-relaxed_rank3 | 6.33 | 7.002 |
| **16** | relaxed_rank1 | 3.597 | 3.842 | 16 | relaxed_rank1 | 3.25 | 6.16 |
| **16** | relaxed_rank3 | 3.43 | 3.618 | 16 | relaxed_rank2 | 0.725 | 1.78 |
| **16** | relaxed_rank4 | 3.789 | 3.516 | 16 | relaxed_rank3 | 2.986 | 2.956 |
| **16** | relaxed_rank5 | 2.385 | 2.478 | 16 | relaxed_rank5 | 3.33 | 3.597 |
| **16** | not-relaxed_rank1 | 1.392 | 3.246 | 16 | not-relaxed_rank1 | 3.207 | 7.093 |
| **16** | not-relaxed_rank2 | 0.715 | 7.719 | 16 | not-relaxed_rank2 | 0.729 | 1.782 |
| **16** | not-relaxed_rank3 | 7.831 | 7.812 | 16 | not-relaxed_rank3 | 5.773 | 7.242 |
| **16** | not-relaxed_rank4 | 1.048 | 7.701 | 16 | not-relaxed_rank4 | 2.98 | 4.657 |
| **16** | not-relaxed_rank5 | 1.307 | 2.539 | 16 | not-relaxed_rank5 | 0.573 | 0.894 |
| **32** | relaxed_rank1 | 3.59 | 3.788 | 32 | relaxed_rank1 | 3.251 | 6.452 |
| **32** | relaxed_rank3 | 5.939 | 8.139 | 32 | relaxed_rank2 | 0.714 | 1.785 |
| **32** | relaxed_rank4 | 3.812 | 3.516 | 32 | relaxed_rank3 | 2.17 | 1.72 |
| **32** | relaxed_rank5 | 2.37 | 2.455 | 32 | relaxed_rank4 | 2.9 | 3.113 |
| **32** | not-relaxed_rank1 | 2.352 | 2.409 | 32 | not-relaxed_rank1 | 0.941 | 1.691 |
| **32** | not-relaxed_rank2 | 0.727 | 7.142 | 32 | not-relaxed_rank3 | 1.475 | 1.707 |
| **32** | not-relaxed_rank3 | 7.341 | 7.998 | 32 | not-relaxed_rank5 | 3.506 | 0.879 |
| **32** | not-relaxed_rank4 | 3.774 | 3.539 | 64 | relaxed_rank1 | 3.256 | 6.188 |
| **64** | relaxed_rank1 | 3.614 | 3.785 | 64 | relaxed_rank2 | 0.719 | 1.775 |
| **64** | relaxed_rank3 | 3.718 | 6.518 | 64 | relaxed_rank3 | 1.537 | 7.436 |
| **64** | relaxed_rank4 | 3.793 | 3.523 | 64 | relaxed_rank4 | 5.98 | 8.142 |
| **64** | relaxed_rank5 | 2.422 | 2.468 | 64 | relaxed_rank5 | 3.202 | 6.326 |
| **64** | not-relaxed_rank1 | 1.394 | 2.98 | 64 | not-relaxed_rank2 | 0.724 | 1.034 |
| **64** | not-relaxed_rank2 | 0.732 | 7.17 | 64 | not-relaxed_rank4 | 2.966 | 8.896 |
| **64** | not-relaxed_rank3 | 6.051 | 8.4 | 8 | #relaxed_rank5 | 0.6 | 7.078 |

| | | | | | | | |
|---|---|---|---|---|---|---|---|
| **64** | not-relaxed_rank4 | 3.241 | 7.717 | 8 | not-relaxed_rank1 | 0.978 | 7.593 |
| **8** | not-relaxed_rank1 | 1.399 | 1.945 | 8 | not-relaxed_rank4 | 1.457 | 7.816 |
| **8** | not-relaxed_rank4 | 1.163 | 1.776 | 16 | relaxed_rank4 | 1.166 | 7.965 |
| **8** | not-relaxed_rank5 | 1.298 | 1.625 | 32 | #relaxed_rank5 | 1.009 | 7.781 |
| **16** | #relaxed_rank2 | 0.758 | 1.804 | 32 | not-relaxed_rank2 | 0.715 | 7.71 |
| **32** | #relaxed_rank2 | 0.739 | 1.763 | 32 | not-relaxed_rank4 | 1.062 | 8.795 |
| **32** | not-relaxed_rank5 | 1.304 | 1.331 | 64 | not-relaxed_rank1 | 0.919 | 7.218 |
| **64** | #relaxed_rank2 | 0.755 | 1.806 | 64 | not-relaxed_rank3 | 1.455 | 6.686 |
| **64** | not-relaxed_rank5 | 1.086 | 1.33 | 64 | not-relaxed_rank5 | 0.579 | 7.264 |
